# Non-helical *Helicobacter pylori* show altered gland colonization and elicit less gastric pathology during chronic infection

**DOI:** 10.1101/505107

**Authors:** Laura E. Martínez, Valerie P. O’Brien, Christina K. Leverich, Sue E. Knoblaugh, Nina R. Salama

## Abstract

Half of all humans harbor *Helicobacter pylori* in their stomachs. Helical cell shape is thought to facilitate *H. pylori’s* ability to bore into the protective mucus layer in a corkscrew-like motion, thus enhancing colonization of the stomach. *H. pylori* cell shape mutants show impaired colonization of the mouse stomach, highlighting the importance of cell shape in infection. To gain a deeper understanding of how helical cell morphology promotes host colonization by *H. pylori*, we used 3D-confocal microscopy to visualize the clinical isolate PMSS1 and an isogenic straight rod mutant (Δ*csd6*) within thick longitudinal mouse stomach sections and performed volumetric image analysis to quantify the number of bacteria residing within corpus and antral glands in addition to measuring total colony forming units (CFU). We found that straight rods show attenuation during acute colonization of the stomach (one day or one week post-infection) as measured by total CFU. Our quantitative imaging revealed that wild-type bacteria extensively colonized antral glands at one week post-infection, while *csd6* mutants showed variable colonization of the antrum at this timepoint. During chronic infection (one or three months post-infection), total CFU were highly variable, but similar for wild-type and straight rods. Both wild-type and straight rods persisted and expanded in corpus glands during chronic infection. However, the straight rods showed reduced inflammation and disease progression. Thus, helical cell shape contributes to tissue interactions that promote inflammation during chronic infection, in addition to facilitating niche acquisition during acute infection.

## Introduction

*Helicobacter pylori* is a Gram-negative, helical shaped bacterium that has evolved to survive in the human stomach. *H. pylori* chronically colonizes the gastric mucosa of approximately 20% of the population in developed countries and greater than 70% of the population in the developing world (1). Most *H. pylori* infections are asymptomatic; however, chronic infection increases the risk of developing chronic active gastritis, peptic ulcer disease, duodenal ulcers, gastric adenocarcinoma, and gastric extranodal marginal zone lymphoma of mucosa-associated lymphoid tissue type (MALT lymphoma) (2). The stomach is an unfavorable environment for bacteria due to its acidity, active digestive enzymes, and low partial oxygen pressure. As a neutrophile, *H. pylori* can only survive minutes in the stomach lumen and overcomes the acidic environment using urease, an enzyme that hydrolyzes urea to produce NH3, locally elevating the pH to near neutral. Successful colonization of the stomach by *H. pylori* requires both urease (3–6) and flagellar-mediated, chemosensory-directed motility to swim out of the lumen and through the mucus layer (7–10). Helical cell shape is thought to facilitate *H. pylori’s* ability to bore into the mucus layer in a corkscrew-like motion, further enhancing its motility through the highly viscous gastric mucus layer that overlies the gastric epithelium. Upon penetrating this thick (~300 μm) mucus layer, *H. pylori* preferentially colonizes a narrow band (~25 μm thick) of mucus immediately overlying the gastric epithelial cell surface (11). While *H. pylori* actively adheres to gastric epithelial cells, it remains extracellular and is only rarely observed within cells (12–14). Upon attachment, *H. pylori* disrupts the tight junctions of epithelial cells to exploit them as a site for replication, where the bacteria can grow as cell-associated microcolonies (15). *H. pylori* also penetrates gastric pits and grows in microcolonies deep in the gastric glands (8).

The helical cell shape of *H. pylori* is generated and maintained by the peptidoglycan (PG) cell wall (16, 17). PG modifying enzymes (Csd1, Csd2, Csd3/HdpA, Csd4, and Csd6) are required to maintain helical cell shape in *H. pylori* (16–22). We and others have shown that *H. pylori* cell shape mutants (Δ*csd1* curved rod and Δ*csd3* variably “c”-shaped rod mutants) are attenuated in stomach colonization (16, 19). Straight rod mutants (Δ*csd4*) also show reduced stomach colonization loads and are outcompeted by wild-type bacteria during co-infection (17). The cell shape mutants do not show a defect in their ability to infect human gastric adenocarcinoma (AGS) cells *in vitro* or to release the pro-inflammatory cytokine interleukin-8 (IL-8) (17). We previously showed that variation in both cell body helical parameters (helical pitch and radius) and flagellum number among different *H. pylori* clinical isolates (LSH100, PMSS1, and B128) leads to distinct and broad swimming speed distributions that reflect both temporal variation in the swimming speed of individual bacterial cells and morphologic variation within the population (23). Furthermore, isogenic mutants with straight rod morphology (Δ*csd6*) showed reduced swimming speeds and a higher fraction of immobilized bacteria in purified gastric mucin gels (23). Whether altered motility behavior in mucus fully accounts for the altered stomach colonization potential of cell shape mutants remains an open question.

Different *H. pylori* strains preferentially colonize the corpus or antrum of the human stomach. The inner lining of the stomach consists of four layers, the serosa, muscularis, submucosa, and mucosa. The mucosa is densely packed with branched tubular gastric glands. Corpus glands are mostly comprised of chief cells, which secrete pepsin, and parietal cells, which secrete hydrochloric acid. The antrum, which comprises about one-fourth of the stomach, is lined by glands mostly containing mucus secreting cells and endocrine cells. Chronic infection with *H. pylori* triggers inflammation in the corpus or antrum, further resulting in distinct disease outcomes. Antral-predominant gastritis is associated with increased acid production, a risk factor for duodenal ulcers (24, 25). Corpus-predominant gastritis leads to loss of parietal cells and eventual reduced acid secretion, increasing the risk for gastric cancer (24, 26, 27). As in the human stomach, *H. pylori* can colonize both the corpus and antrum regions of the mouse gastric mucosa (28). The mouse stomach contains two grossly distinct stomach regions, a non-glandular (forestomach) region and a glandular region (corpus and antrum). The forestomach, which does not become colonized by *H. pylori*, is lined with keratinized squamous epithelium and is separated from the glandular corpus region by a raised mucosal fold referred to as the limiting ridge. Several mouse-adapted *H. pylori* isolates induce gastritis and gland atrophy in C57BL/6 mice, but do not induce neoplasia (29, 30). However, chronic infection with PMSS1, a strain shown to be more virulent than other clinical isolates in mice, triggers inflammation, gland hyperplasia, gastric atrophy, and early signs of metaplasia (31, 32).

Here, we used a mouse model of infection to investigate how helical cell shape helps *H. pylori* establish infection and acquire a replicative niche within the stomach. In addition to enumerating colony forming units (CFU) of *H. pylori* in homogenized stomach tissue, we used 3D-confocal microscopy and volumetric image analysis to localize and quantify the number of bacteria present within corpus and antral glands. We discovered that Δ*csd6* straight rods are attenuated at one day and one week post-infection, both in CFU load and localization within gastric glands, yet can nonetheless establish chronic infection. At one week post-infection, straight rods show reduced localization within antral glands, while at one month post-infection, localization within both antral and corpus glands is similar to or greater than that of wild-type *H. pylori*. In spite of their ability to localize within the glands, Δ*csd6* straight rods elicited less inflammation and hyperplasia in the antrum and the transition zone between corpus and antrum at one and three months post-infection. Our study supports a role for helical cell shape in promoting efficient stomach colonization during acute infection and in driving gastric pathology during chronic infection.

## Results

### Csd6-dependent helical cell shape of *H. pylori* confers an advantage during initial colonization of the stomach

To investigate how helical cell morphology contributes to *H. pylori* stomach colonization and persistence, we conducted single-strain infections and competitions with wild-type *H. pylori* PMSS1 and an isogenic straight rod mutant (Δ*csd6*, Table S1). We harvested a third of the stomach to assess bacterial load by colony-forming units (CFU) per gram of stomach tissue, a third to fix in paraformaldehyde for immunohistochemistry for enumeration of bacteria within gastric glands, and a third for pathological evaluation (Fig. 1). As expected, we did not recover *H. pylori* from the mock-infected groups at any time point (data not shown). The Δ*csd6* straight rod mutant showed significantly attenuated gastric loads at one day with a two log difference in CFU/g of stomach tissue compared to wild-type (*p* < 0.0001, unpaired non-parametric two-tailed Mann Whitney U-test) (Fig. 2A). At one week, the mutant had a one log difference in recovery compared to wild-type (*p* < 0.0001), as had been previously reported for another straight rod mutant *(Δcsd4)* generated in a different *H. pylori* strain background (LSH100) (17). In a competition experiment, wild-type bacteria strongly outcompeted the Δ*csd6* mutant at one week post-infection (*p* = 0.0005, paired t-test) (Fig. 2B). Complementation of the Δ*csd6* mutant by expressing the *csd6* gene at a distal intragenic locus (22, 33), restored helical cell shape and no significant differences in side curvature distributions were observed between wild-type and the *csd6* complemented strain (*p* = 0.6, Kolmogorov-Smirnov statistics) (Fig. S1). In mice, the *csd6* complemented strain showed comparable colonization loads to the wild-type strain at one day post-infection, but interestingly was comparable to the Δ*csd6* mutant at one week post-infection (Fig. 2A). However, in competitive infection, the Δ*csd6* mutant was outcompeted by the *csd6* complemented strain in 6 of 10 mice at one week in two independent experiments (Fig. 2B). Taken together, these data suggest that the *csd6* complemented strain has some shape-independent colonization defect that manifests after the first day of infection. Nonetheless, we tested both the deletion strain and the complemented strain in subsequent experiments.

**Figure 1.**
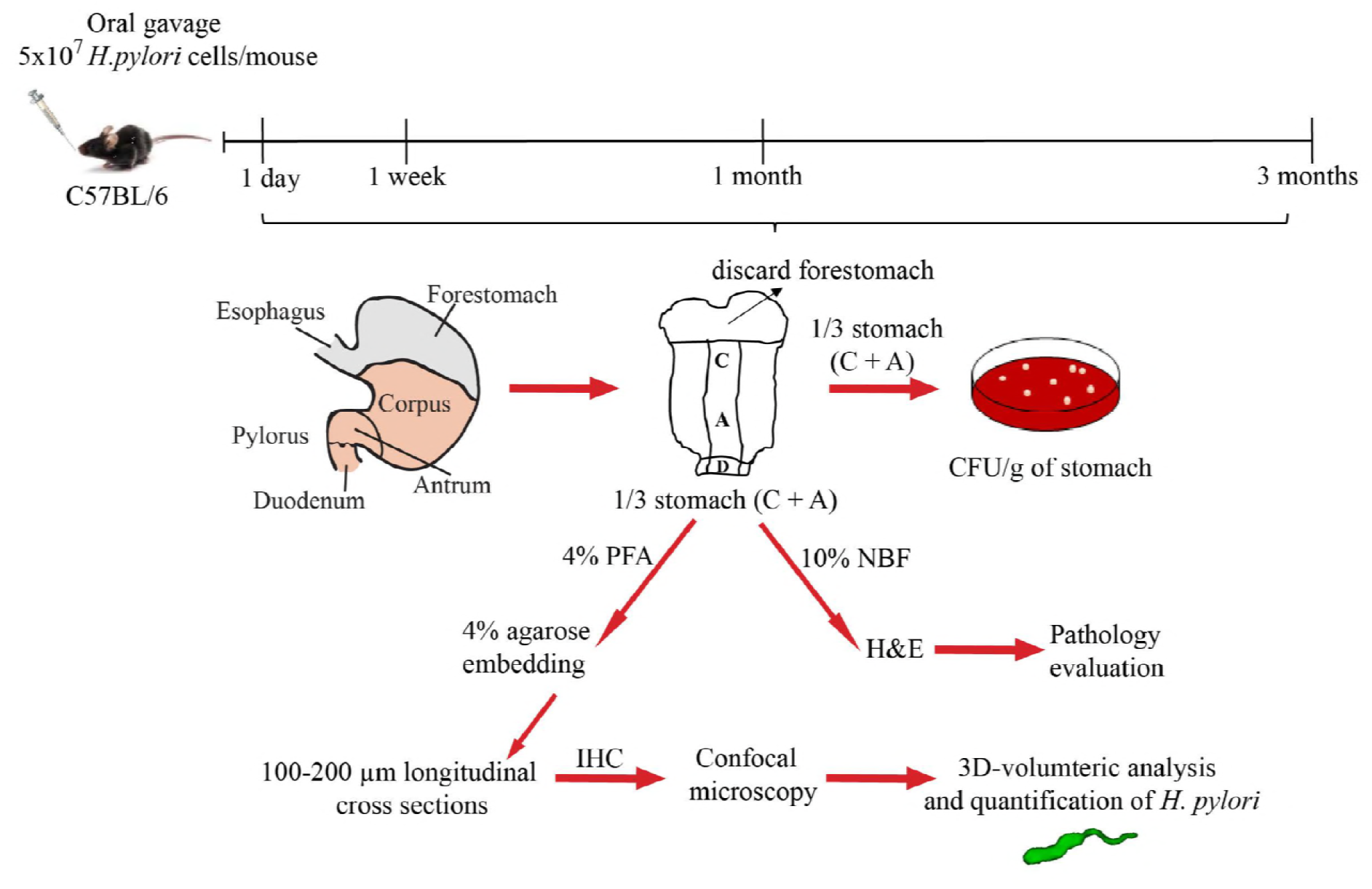
Experimental outline. C57BL6/J mice were infected by oral gavage with wild-type (PMSS1), straight rod mutants (Δ*csd6*), or *csd6* complemented *H. pylori* bacteria, or mock-infected with broth. At the indicated time points, the stomach was removed and one third used to determine bacterial load, one third for pathology evaluation, and one third for bacterial localization within glands. C, corpus; A, antrum; CFU, colony-forming units; PFA, paraformaldehyde; NBF, neutral-buffered formalin; H&E, hematoxylin and eosin; IHC, immunohistochemistry.

**Figure 2.**
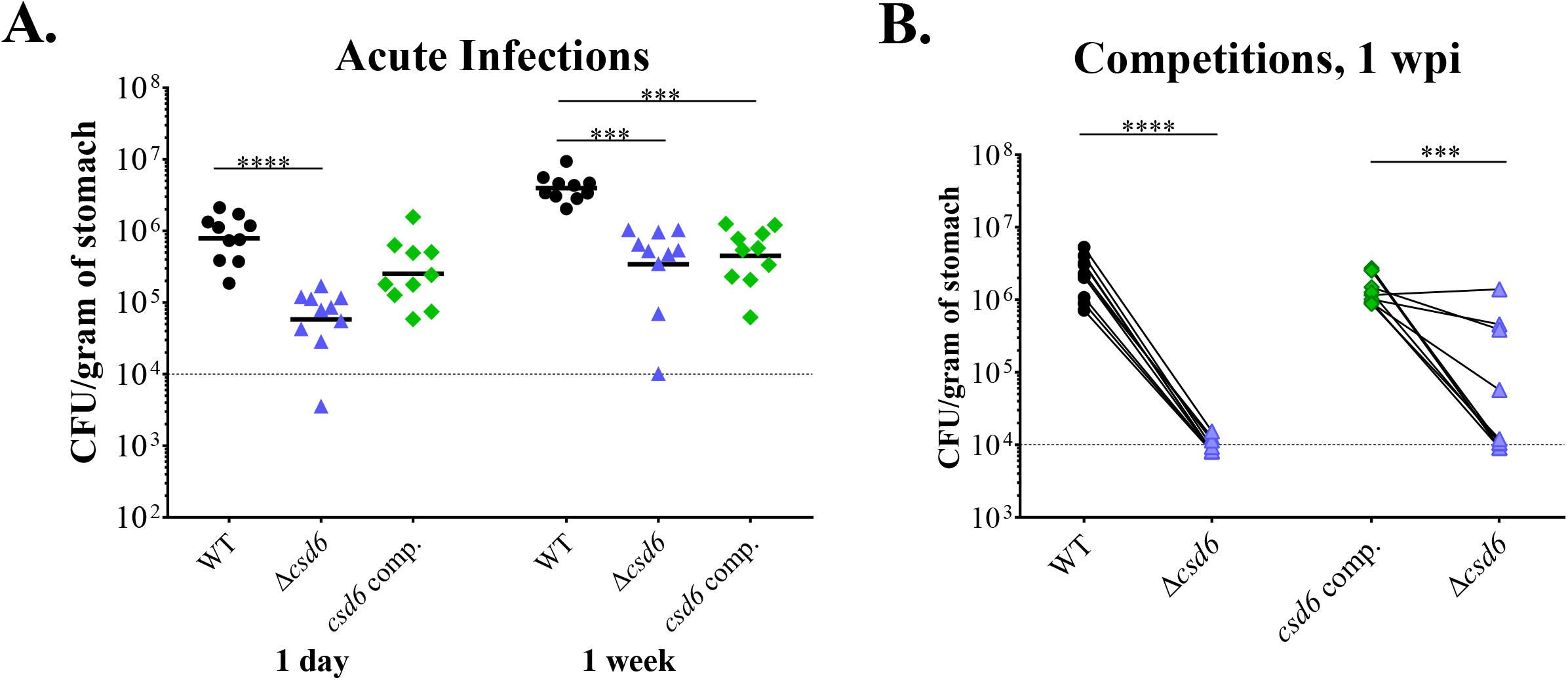
The *H. pylori* straight rod mutant Δ*csd6* shows early colonization defects compared to wild-type bacteria. Single or competitive infections were performed with the wild-type strain, Δ*csd6, csd6* complemented, or broth (mock-infection control). (**A**) Stomach loads at one day and one week of infection. *** *P* <0.001, **** *P* < 0.0001, Kruskal-Wallis test with Dunn’s multiple test correction. (**B**) Competitive infections between wild-type and *Δcsd6*, or *csd6* complemented and *Δcsd6*, with lines connecting the bacterial load values for each genotype from the same mouse. *** *P* < 0.001, **** *P* < 0.0001, Mann-Whitney U test. Dotted line indicates the average limit of detection. Data are from two independent experiments with n=10 mice per group. WT, wildtype; comp, complemented; wpi, week post-infection.

**Table 1.** Bacterial strains used in this study.

| Strain Name | Genotype or Description | Shape phenotype | Reference or Source |
| --- | --- | --- | --- |
| PMSS1 | Wild-type *H. pylori* | helical | (31, 45) |
| LMH6 | $\Delta csd6::cat$ in PMSS1 | straight | This study |
| TSH17 | $\Delta csd6::cat$ in LSH100 | straight | (22) |
| TSH31 | $\Delta csd6::cat$ McGee:*csd6:aphA3* in LSH100 | helical and straight morphologies | (22) |
| EPH1 | *csd6* complemented strain: $\Delta csd6::cat$ McGee:*csd6:aphA3* in PMSS1 | helical | This study |
| LMH12 | *csd6* complemented strain: EPH1 backcrossed in PMSS1 | helical | This study |

### 3D-volumetric image analysis of *H. pylori* indicates an antral gland preference for both wild-type and straight rod mutant bacteria during acute infection

Others have shown that during experimental infection in C57BL/6 mice, *H. pylori* bacteria reside in the mucus layer that overlies the stomach epithelium and a subpopulation of bacteria penetrate deep in the gastric glands, where they can adhere to gastric epithelial cells comprising the mid-glandular proliferative zone (8, 32). We questioned whether the Δ*csd6* mutant could occupy the same niches. In thin (4-5 μm) sections, *H. pylori* is difficult to quantify, because the gastric gland lumen is rarely captured. Thus, we followed a recently established method for 3D-confocal microscopy to visualize and quantify the number of *H. pylori* bacteria colonizing corpus and antral glands in thick (100-200 μm) sections using volumetric image analysis of individual bacterial cells and bacteria within microcolonies (8, 32). Similar to prior studies, the mean volume for individual cells analyzed was 9.5 μm^3^ (n = 203 cells analyzed) and clusters of two bacteria showed a proportional increase in volume (Fig. S2).

At one day post-infection, both wild-type and the Δ*csd6* mutant were detected in corpus and antral glands at very low densities (data not shown). At one week, 3D-visualization of the antrum showed wild-type bacteria associated with gastric epithelial cells near the luminal surface, as well as deeper in the glands, where they form dense microcolonies (Fig. 3A). To explore bacterial localization differences among strains, we quantified the number of bacteria in each field of view along the length of the stomach for one mouse from each genotype with similar colonization loads (Fig. 3B-D; wild-type 7.6×10^5^ CFU/g, Δ*csd6* 4.0×10^5^ CFU/g, *csd6* comp. 5.4×10^5^ CFU/g). Wild-type bacteria were easily detected in antral glands and the transition zone between the corpus and antrum (C/A) junction (Fig. 3B). Fewer bacteria were observed in corpus glands (<40 bacteria per field of view). The Δ*csd6* mutant bacteria were detectible but at lower numbers in both corpus and antral glands (fewer bacteria per gland as well as fewer glands colonized) (Fig. 3C and S3A). Like wild-type, *csd6* complemented bacteria were observed predominantly in the antrum (Fig. 3D and S3B) and the C/A junction. We extended this analysis to additional animals from each genotype (Fig. 3E). In all animals infected with wild-type and *csd6* complemented strains, as well as two out of three animals infected with *Δcsd6*, the bacteria preferentially localized to the antrum instead of the corpus. However, the levels of bacteria detected in the glands did not correlate well with CFU loads. While there was a trend toward lower levels of gland localization for Δ*csd6* compared to wild-type and *csd6* complemented bacteria, the Δ*csd6* mutant was able to penetrate and multiply within both corpus and antral glands in at least a subset of animals, despite exhibiting a significantly lower CFU load at this time point.

**Figure 3.**
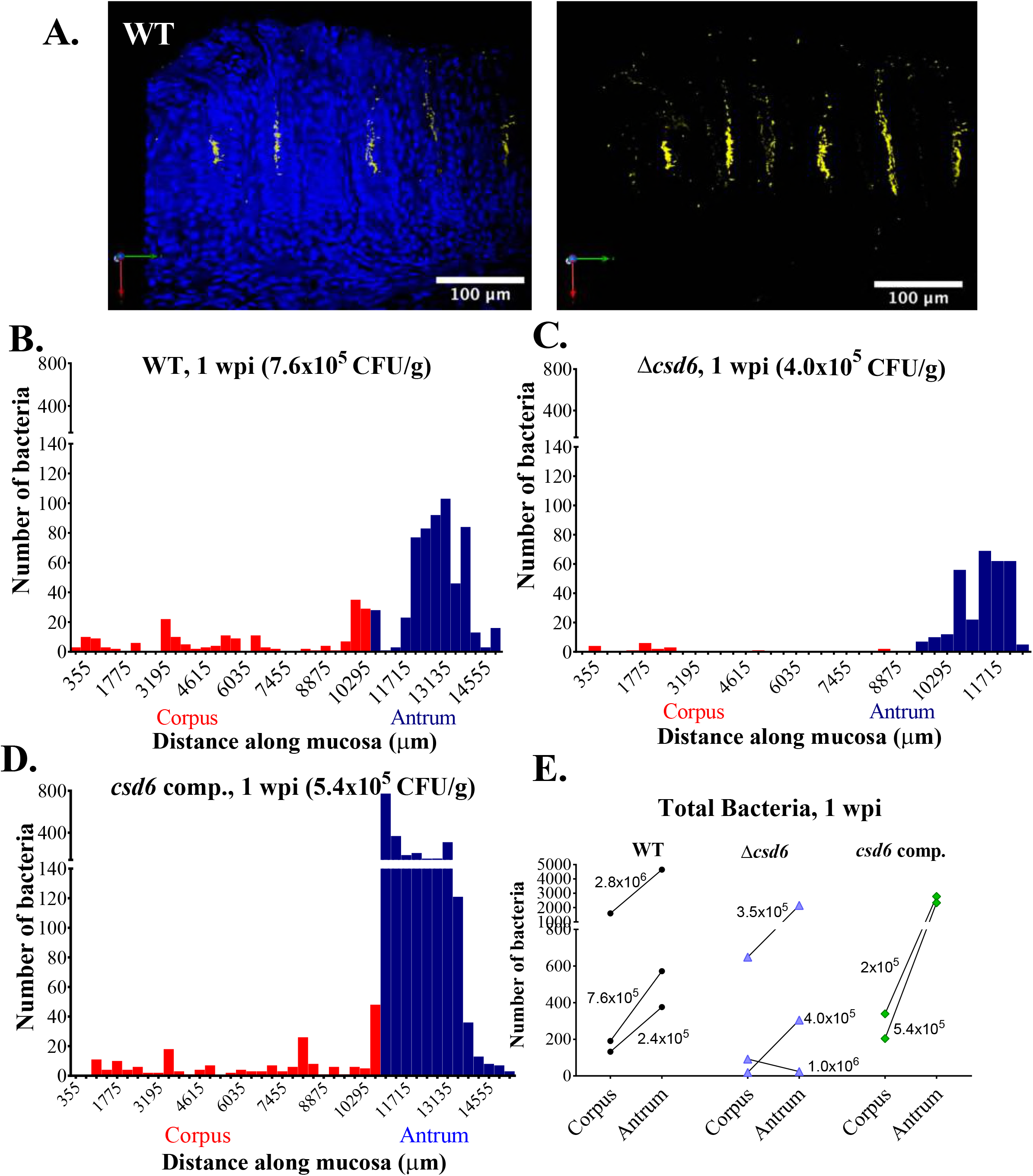
The Δ*csd6* straight rod mutant is attenuated in colonizing the corpus and antrum at one week post-infection. Thick stomach sections from the one week infections shown in **Figure 2A** were stained for *H. pylori* and the number of bacteria within the glands was quantified along the entire length of the section. (**A**) Representative images of the antrum of a mouse infected with wild-type *H. pylori* for one week. Shown are maximum intensity projections of Z-stacks, with blue (DAPI, left panel) staining nuclei and yellow staining *H. pylori*. Scale bar = 100 μm. (**B – D**) Gland analysis for wild-type *H. pylori* (**B**, same mouse as **A**), Δ*csd6* (**C**) and *csd6* complemented (**D**), showing the number of bacteria detected by immunofluorescence within glands along the length of the stomach in microns. Red bars indicate the corpus and blue indicate the antrum. (**E**) The total number of bacteria in the corpus and antral glands is shown for n = 2-3 mice per strain, with the CFU per gram of stomach for each mouse indicated on the graph. WT, wildtype; comp, complemented; wpi, week post-infection.

### Both helical *H. pylori* and straight rods show expansion into the corpus after one month

At one month post-infection, we observed a wide distribution of stomach loads from all strains tested, but in each case the geometric mean was around 10^5^ CFU/g (Fig. 4A). A subset of mice showed stomach loads of 10^6^ CFU/g or more, while other mice showed low loads near or below the limit of detection (10^3^ CFU/g). At three months, stomach loads from each infected group were more tightly clustered, though the geometric mean was still around 10^5^ CFU/g (Fig. 4A). Thus, Csd6-mediated helical cell shape is necessary for robust stomach colonization during the acute stages of gastric infection in mice, but not for maintenance of chronic infection.

**Figure 4.**
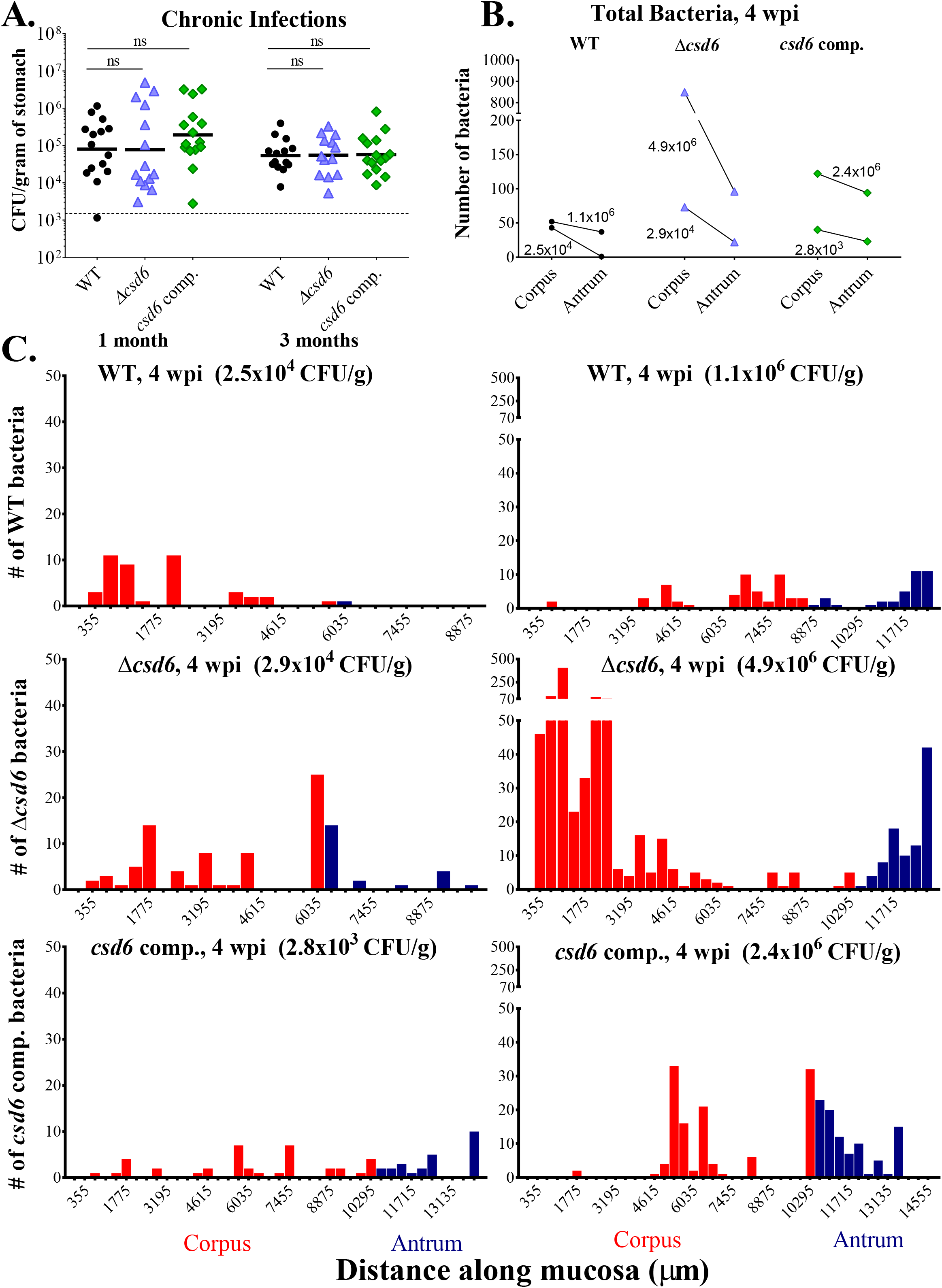
Both wild-type *H. pylori* and the Δ*csd6* straight rod mutant can expand into the corpus by one month post-infection. Single infections were performed with the wild-type strain, Δ*csd6, csd6* complemented, or broth (mock-infection control). (**A**) Stomach loads at one month and three months post-infection. ns, not significant by Kruskal-Wallis test with Dunn’s multiple test correction. Data are from two independent experiments with n=15 mice per group; the limit of detection is shown with a dotted line. (**B** and **C**) Thick stomach sections from the one month infections shown in A were stained for *H. pylori* and the number of bacteria within the glands was quantified along the entire length of the section. For each bacterial strain, a mouse with a “low” CFU load (left panel) and a “high” CFU load (right panel) was analyzed. (**B**) The total number of bacteria in the corpus and antrum is shown for n = 2 mice per bacterial strain, with the CFU per gram of stomach for each mouse indicated on the graph. (**C**) Gland analysis for wild-type *H. pylori, Δcsd6* and *csd6* complemented strains, showing the number of bacteria within corpus and antral glands along the length of the stomach in microns. Red bars indicate the corpus and blue indicate the antrum. WT, wildtype; comp, complemented; wpi, week post-infection.

Given the variable stomach CFUs at one month (Fig. 4A), we determined the localization of bacteria in gastric tissue samples with either “low” (~10^4^ CFU/g of stomach) or “high” (~10^6^ CFU/g of stomach) bacterial loads of the three strains (Fig. 4B). We observed two key differences from the one week analysis (compare to Fig. 3E): first, the total number of bacteria quantified at one month was generally lower than at one week; and second, at one month there were more bacteria in the corpus than the antrum – the reverse of what was observed at one week. For mice with a “low” CFU load (Fig. 4C, left panels), the numbers of bacteria detected in corpus glands were similar among bacterial genotypes (fewer than 75 total bacteria). However, both the Δ*csd6* and the *csd6* complemented strains had more bacteria in the antrum (22 and 23 total bacteria, respectively) than the wild-type strain did (one bacterium). For mice with a “high” CFU load (Fig. 4C, right panels), the Δ*csd6* mutant was different from the other two strains, with more bacteria detected in the corpus (849) than the antrum (96). The wild-type and *csd6* complemented strains had more similar numbers of bacteria in the corpus (52 and 122, respectively) and antrum (37 and 94, respectively). While we did detect more total bacteria in the “high” CFU mice than their “low” CFU counterparts, CFU load did not correlate well with the number of bacteria detected in the glands.

### Chronic infection with the straight rod mutant results in reduced inflammation

Next, we assessed pathologic responses in mice infected with the different genotypes. The most severe lesions in each animal were scored according to previously developed pathology scoring criteria for inflammation, epithelial defects, oxyntic atrophy, hyperplasia, and metaplasia in the corpus and inflammation and hyperplasia in the antrum (Table S2 and (34)). Individual scores (0-4) for each criterion were summed to generate a histological activity index (HAI) score. At one and three months post-infection, animals showed evidence of both inflammation and hyperplasia throughout the stomach. As shown in Fig. 5, which show representative images of animals with the highest HAI score for each genotype, pathologic changes were most pronounced at the corpus/antrum junction (C/A). Oxyntic atrophy (loss of parietal cells) and metaplasia were also observed within the distal corpus near the C/A junction (e.g. Fig. 5B).

**Figure 5.**
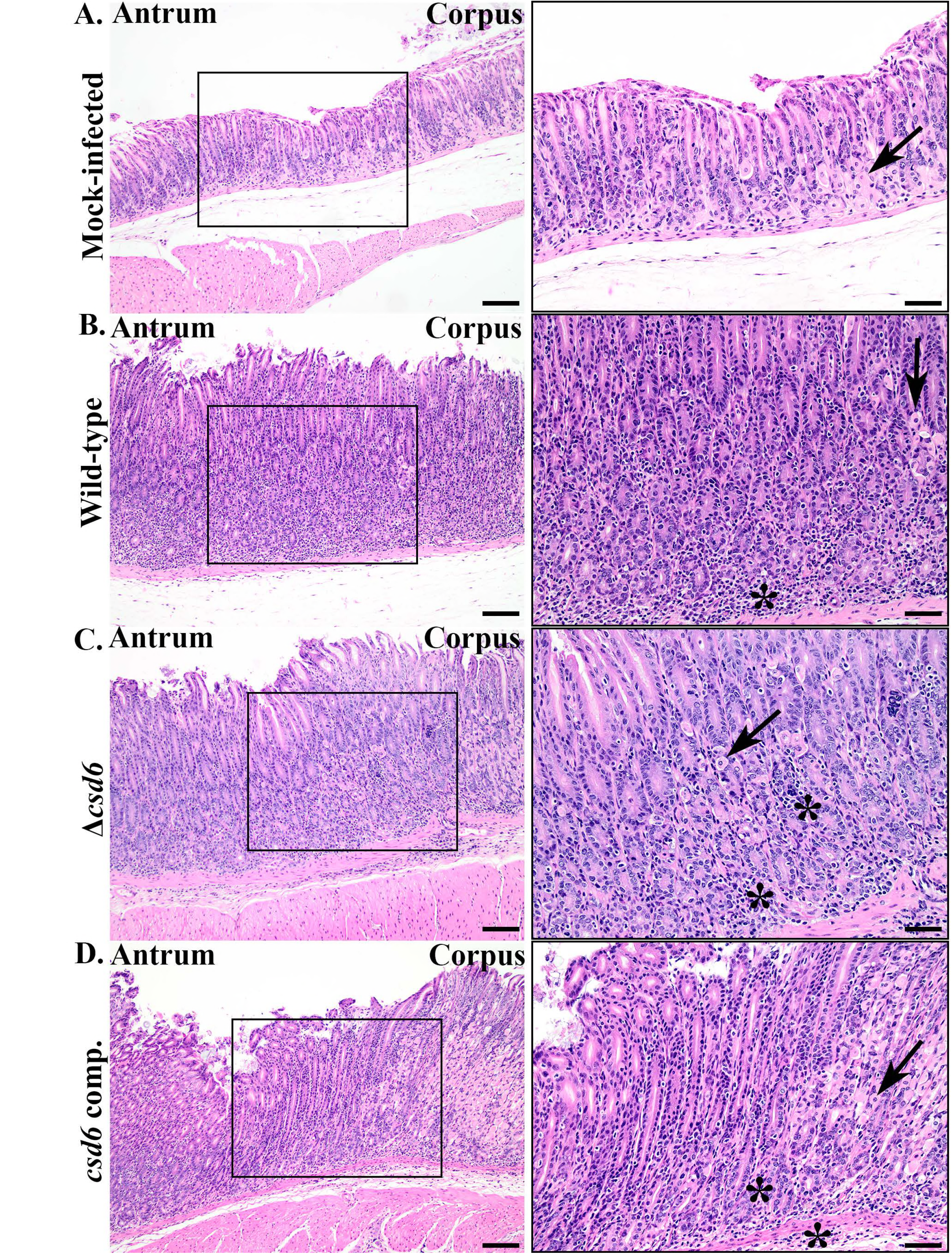
The Δ*csd6* straight rod mutants elicit less immunopathology compared to wild-type and *csd6* complemented bacteria. Images of hematoxylin and eosin-stained sections from the three month infection, showing corpus, corpus/antral junction (box), and antral regions of the most severe histopathologic changes in each group. Right panels show higher magnification images (20x) within the enclosed black boxes of 10x images on the left. Arrows point to remaining parietal cells in corpus glands and the asterisks denote sites of infiltrating inflammatory cells. Images are from (**A**) Mock-infected, (**B**) wild-type-infected (HAI = 21, 7.8 x10^3^ CFU/g stomach), (**C**) Δ*csd6*-infected (HAI = 15, 1.2 x10^5^ CFU/g stomach), and (**D**) *csd6* complemented-infected (HAI = 22, 4.0 x10^4^ CFU/g of stomach) mice. WT, wildtype; comp, complemented; HAI, histological activity index. Left panels scale bar =100 μm; right panels scale bar = 50 μm.

At one month post-infection, wild-type and *csd6* complemented bacteria induced inflammation characterized by lymphocytic and neutrophilic infiltrates and low level oxyntic atrophy at the antrum and C/A junction. In contrast, mice infected with the Δ*csd6* mutant had little to no gastric inflammation and reduced hyperplasia (Fig. 6A and S4A-B), even though about half the mice had fairly high bacterial loads and bacteria were detected within gastric glands at this time point (Fig. 4A). The differences in pathology among bacterial genotypes became more pronounced at three months (Fig. 6A and S4C-D), by which time the Δ*csd6* mutant infected animals showed significantly reduced HAI scores and individual scores for inflammation, oxyntic atrophy, and hyperplasia in the C/A junction (Fig. S4C-D). Animals infected with wild-type bacteria showed evidence of an inverse relationship between stomach CFU load and HAI (Spearman correlation coefficient r = −0.22, Fig. 6B), which is consistent with prior studies showing lower loads in animals with more severe gastritis (35). In contrast, animals infected with the Δ*csd6* mutant showed the opposite trend (Spearman correlation coefficient r = 0.518, Fig. 6B). Thus, although bacterial loads are not statistically significantly different at three months post-infection (Fig. 4A), the Δ*csd6* mutant elicits significantly less inflammation and hyperplasia compared to wild-type or the *csd6* complemented strain at this time point.

**Figure 6.**
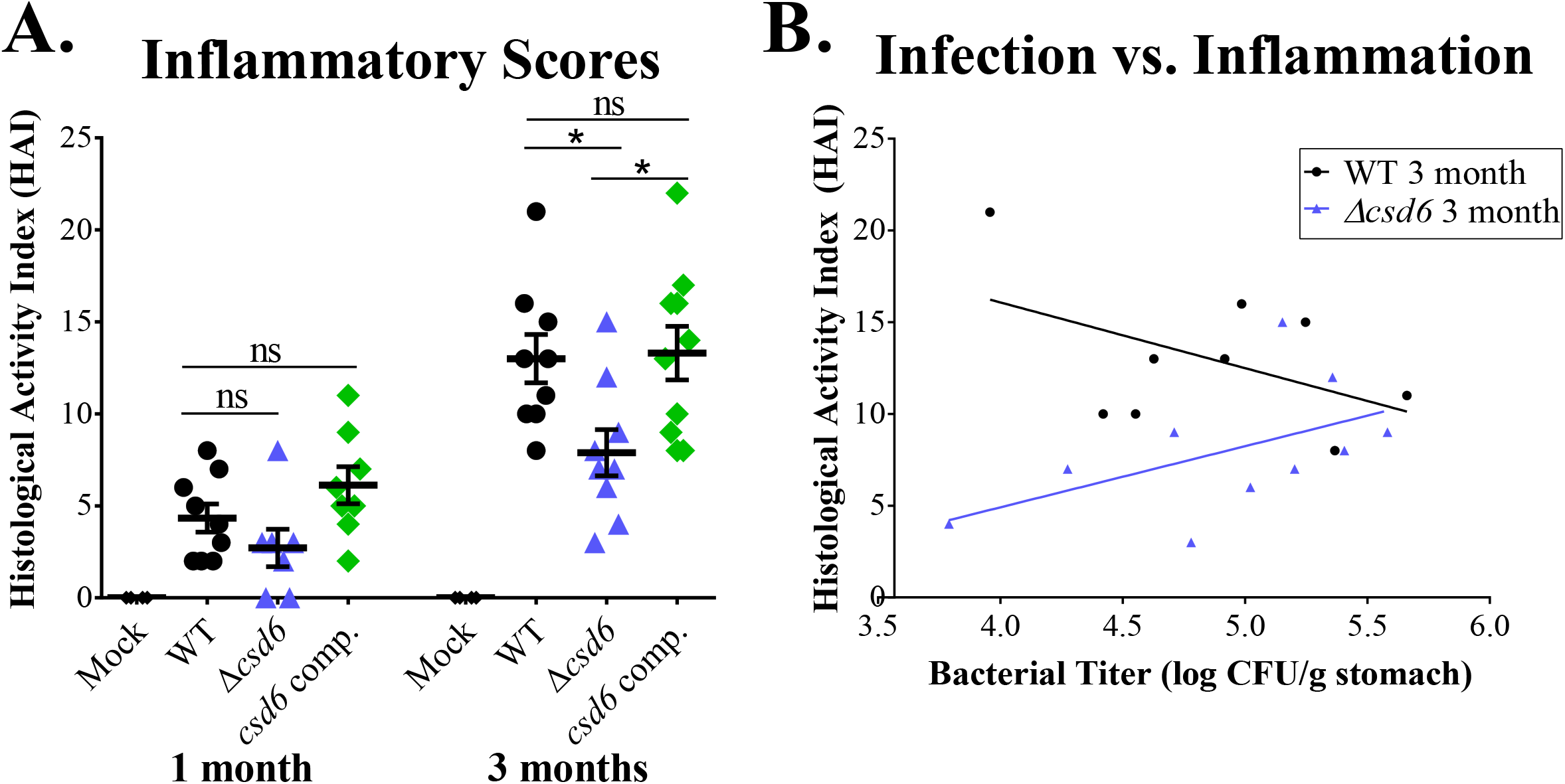
Chronic Δ*csd6* mutant infections show significantly less histological activity compared to wild-type and *csd6* complemented infections. Thin stomach sections from the mice in Figure 4A were used for a blinded analysis of stomach inflammation and pathology. (**A**) The total histological activity index (HAI) is provided for mock-infected (“Mock”), wild-type, Δ*csd6*, and the *csd6* complemented strain at one and three months of infection. Mean ± standard deviations are shown. * *P* <0.05, Kruskal-Wallis test with Dunn’s multiple test correction. (**B**) Plot showing the correlation between wild-type and Δ*csd6* stomach colonization loads and total HAI.

## Discussion

It has long been proposed that helical cell shape facilitates *H. pylori*’s ability to penetrate the thick gastric mucus layer by enhancing cell body propulsion, thus promoting colonization of the stomach (36). Our study confirms and extends prior results that helical cell shape, while not required for stomach colonization, confers a significant advantage to *H. pylori* during acute infection (one day and one week) (16, 17). During chronic infection, the colonization levels of the wild-type strain and the Δ*csd6* mutant were comparable, suggesting that the mutant can nonetheless chronically colonize glands of the stomach. Thus, straight rods may occupy a specific niche within the stomach allowing them to persist long-term. By performing 3D image analysis of *H. pylori* in thick stomach sections we found that wild-type and *csd6* complemented *H. pylori* had reduced localization within corpus and antral glands at one month post-infection relative to the number of bacteria observed in the glands at one week, which may be attributed to adaptive immune responses clearing the infection (35). While Δ*csd6* infected tissues appeared to have somewhat lower levels of bacteria within the glands at one week, the contraction of the gland population at one month appeared less dramatic, particularly in the corpus. Our fixation conditions do not preserve the mucus layer overlying the epithelium, which may host a significant fraction of bacteria in both mutant and wild-type infections. Finally, we found that the Δ*csd6* mutant elicited less inflammation than wild-type or complemented bacteria at one and three months post-infection, despite having comparable bacterial CFU loads and similar or greater localization within the gastric glands. Thus, our study is the first to suggest a bacterial load-independent link between *H. pylori*’s helical cell shape and chronic gastritis.

During chronic infection with *H. pylori*, the degree of inflammation found at the corpus/antrum junction is often greater than observed in the adjacent mucosa of either the corpus or antrum, and may promote glandular atrophy, loss of parietal cells in the corpus, and eventually mucous cell gland hyperplasia (37). In the present study, we found highly variable colonization loads in mice infected with either wild-type, the Δ*csd6* mutant, or the *csd6* complemented strain for one month, which may be related to differential host responses among mice. Chronic infection with wild-type PMSS1 or the *csd6* complemented strain induced pathology in the antrum and the C/A junction. However, straight rods elicited less inflammation and gland hyperplasia. Chronic infection with *H. pylori* is controlled by innate and adaptive immune responses, regulated by CD4^+^ T-helper 1 (Th1), Th17-polarized T-effector cell subsets, B-cells, and their secreted cytokines (35, 38). Evaluating the adaptive immune response to infection with the Δ*csd6* mutant will determine if its differential localization in corpus glands at one month influenced its ability to induce inflammation or cause gastric disease. In addition, some *H. pylori* strains, including PMSS1 used in this study, express VacA, a virulence factor that has been shown to suppress T-cell responses to mediate longevity of infection (39–41). It will also be important to determine whether *H. pylori* cell shape mutants have unexpected effects on the expression of other factors that could mediate or suppress inflammation, such as VacA, to maintain a favorable niche in the stomach.

Our work demonstrates that the helical cell shape of *H. pylori* is important for acute colonization of the stomach and enhances pathology during chronic infection. Helical cell shape is also important for a bacterial pathogen that colonizes the human intestinal tract, *Campylobacter jejuni*. Isogenic straight rod mutants of *C. jejuni pgp1* and *pgp2* (homologs of *H. pylori csd4* and *csd6*, respectively) show attenuated motility in soft agar (42, 43) similar to *H. pylori*. However, *C. jejuni pgp1* and *pgp2* mutants completely fail to colonize intestinal crypts or to induce inflammatory responses in the mouse model tested (44).

In summary, our study provides insight into how helical cell shape impacts *H. pylori*’s niche acquisition and inflammation in the stomach. Loss of helical cell shape alters *H. pylori*’s ability to utilize some of the available niches within the stomach and its ability to promote inflammation and tissue hyperplasia when present in the glands. As morphological diversity exists between different clinical isolates of *H. pylori* in cell body and helical cell parameters (23), *H. pylori* clinical isolates with decreased helical pitch and twist may differ in their ability to colonize certain gastric niches and in the trajectories of pathogenesis. Thus, diversity in cell shape parameters may contribute to the diversity of pathogenic outcomes observed in infected individuals.

## Materials and Methods

### H. pylori strains and growth conditions

Strains used in this study are described in Table S1. Briefly, wild-type *H. pylori* strain PMSS1, also called 10700 (31, 45), and derivatives were cultured on horse blood plates or in liquid media containing 90% (v/v) Brucella broth (BD Biosciences) and 10% fetal bovine serum (GIBCO) (BB10) in the absence of antimicrobials, as previously described (16). Cells were maintained at 37°C under microaerobic conditions in a trigas incubator equilibrated to 10% CO_2_ and 10% O_2_. Plates were incubated 24-72 hours and liquid cultures were incubated for 12-16 hours under constant agitation at 200 rpm. For resistance marker selection, horse blood plates were supplemented with chloramphenicol (15 μg/mL) or kanamycin (25 μg/mL).

### Strain construction

Isogenic mutant of *csd6* (HPG27_477) in the PMSS1 strain background was generated by transfer of the mutation constructed in the G27/LSH100 strain background (16, 22) using natural transformation (46). Transformants were confirmed by PCR using primers homologous to upstream and downstream flanking regions of the gene using the primers listed in Table S2. The mutation was then backcrossed into PMSS1 once by isolating genomic DNA from the resulting strain for natural transformation of PMSS1. The resulting backcrossed clones were evaluated by PCR to confirm replacement of the wild-type allele with the null allele. Clones were checked for urease activity and motility, and single clones were used for quantitative morphology analysis and for oral gavage of mice.

The *csd6* complemented strain was constructed by natural transformation of PMSS1 Δ*csd6* with genomic DNA from a *csd6* complemented strain generated in the *H. pylori* G27 strain background, TSH31 (Δ*csd6::cat McGee:csd6:aphA3*), where a wild-type copy of *csd6* (HPG27_477) was introduced at a neutral intergenic chromosomal locus (33). Genomic DNA from TSH31 was used for natural transformation of the PMSS1 Δ*csd6* strain. Transformants were selected on horse blood plates supplemented with kanamycin (25 μg/mL). The recipient strain (EPH1) was PCR confirmed using primers homologous to upstream and downstream flanking regions using the primers listed in Table S1. The *csd6* complemented strain was then backcrossed once by isolating genomic DNA from EPH1 for natural transformation of PMSS1 Δ*csd6*. The resulting backcrossed clones were evaluated by PCR to confirm integration of *csd6* and were renamed LMH12 clones 1-3. The clones were then checked for urease activity and motility, and were used for quantitative morphology analysis. Figure S1 shows morphology analysis of LMH12 clone 3 (*csd6* complemented strain), which was the strain used for single-strain infections in mice and for bacterial localization studies.

### Morphology analysis

Wild-type *H. pylori* PMSS1, Δ*csd6*, and *csd6* complemented bacteria were grown in liquid culture to an optical density at 600 nm (O.D. (600)) of 0.4 - 0.6. Bacteria was fixed in 4% Paraformaldehyde with 25% Glycerol in 1X PBS and added to 0.1% Poly-L-lysine (Sigma Aldrich) coated coverslips, which were then placed on a pre-cleaned microscope slide and sealed with VaLP (1:1:1 Vaseline: Lanolin: Paraffin). Single focal plane images were collected using a 100X ELWD Plan APO (NA 1.40 oil) objective mounted on a Nikon TE 200 microscope, equipped with a Nikon CoolSNAP HQ CCD camera controlled by MetaMorph software (MDS Analytical Technologies). Quantitative morphology analysis of manually thresholded phase-contrast images was performed as described in Sycuro *et al*. 2010 using the CellTool software program (16, 47).

### Parameter optimization for H. pylori 3D-image analysis

Wild-type *H. pylori* PMSS1 bacteria was grown in liquid culture to an optical density at 600 nm (O.D. _(600)_) of 0.4 - 0.6. A 1 mL of bacterial culture was harvested in a 1.5 mL microcentrifuge tube and centrifuged at 5,000 rpm for 5 min. The cell pellet was resuspended in 100-200 μL of 4% Paraformaldehyde (PFA) in 100 mM phosphate buffer (pH 7.4) and fixed for at least 30 min at room temperature. Bacteria were then embedded in 4% agarose (ultra-pure low-melting point agarose, Invitrogen) prepared in 1X phosphate-buffered saline (pH 7.4) (Gibco). The agarose solution was first cooled down to ~55 - 65°C and then aliquoted into 1.5 mL microcentrifuge tubes. Aliquots of fixed bacteria were immediately added and gently resuspended into the solution before it solidified. The solidified slabs were gently removed by insertion of a metal spatula on the side of the tube. The slabs were then sectioned using a vibratome (Leica VT 1200 S fully automated vibrating blade microtome, Leica Biosystems, Germany) to generate 100 - 200 μm thick sections. Sections were permeabilized in blocking buffer (3% bovine serum albumin (Sigma Aldrich); 1% Saponin (Sigma Aldrich); and 1% Triton X-100 (Sigma Aldrich) in 1X PBS) and immunostained with primary anti-H. *pylori* rabbit polyclonal antibody (1:1,000 dilution) (gifted by Dr. Manuel Amieva at Stanford University) overnight at 4°C. A goat anti-rabbit Alexa Fluor-488 conjugate antibody (1:2,000) (Molecular Probes) was used to visualize *H. pylori*. Samples were incubated in the secondary antibody for 2 hrs at room temperature. Sections were then washed 5X with blocking buffer and then mounted onto standard glass microscope slides with secure imaging spacers (9 mm diameter x 0.12 mm depth, Electron Microscopy Sciences). Pro-Long Diamond Antifade medium was added (Molecular Probes) before mounting on coverslips.

### Ethics statement

All procedures involving animals were done under practices and procedures of Animal Biosafety Level 2 and carried out with strict accordance with the recommendations in the Guide for the Care and Use of Laboratory Animals of the National Institutes of Health. The facility is fully accredited by the Association for Assessment and Accreditation of Laboratory Animal Care and complies with the United States Department of Agriculture, Public Health Service, Washington State, and local area animal welfare regulations. All activities were approved by the FHCRC Institutional Animal Care and Use Committee (IACUC; protocol number 1531). Animals were euthanized by CO_2_ asphyxiation followed by cervical dislocation.

### Mouse infections

4-6 week old female C57BL/6J mice were purchased from the Jackson Laboratory (Bar Harbor, Maine, U.S.) and were certified free of endogenous *Helicobacter* infection by the vendor. All animals were maintained in autoclaved microisolator cages (1-5 mice per cage) and provided with standard chow and water ad libitum. Mice were infected with a single dose of 5 x 10^7^ *H. pylori* cells/strain (0.1 mL) via oral gavage. Mock-infected controls were gavaged with 0.1 mL of liquid culture media containing 90% (v/v) Brucella broth and 10% fetal bovine serum (BB10); no *H. pylori* were recovered from mock-infected mice. Mice were euthanized by inhalation of CO_2_ and stomachs were harvested at one day, one week, and one or three months post-infection. Most of the non-glandular region (forestomach) was discarded since this region of the stomach is lined with squamous rather than glandular epithelium. *H. pylori* has not been shown to colonize this region of the stomach. However, *H. pylori* may colonize the interface between the squamous forestomach and glandular stomach where the corpus begins (the squamocolumnar junction). Regions of interest for *H. pylori* colonization include the corpus, antrum, and the pyloric junction with the duodenum. Therefore, part of the forestomach was maintained and the glandular stomach (corpus and antrum) was opened along the lesser curvature from the esophagus through the proximal duodenum. For one day and one week harvests, half the stomach was used for plating for CFU enumeration and the other half was fixed in 4% paraformaldehyde (PFA) in 100 mM phosphate buffer (pH 7.4) for 1-2 hours. For chronic infection time points (one and three months), the stomach was divided into thirds. A third of the stomach was collected to measure CFU/gram of stomach load, a third was fixed in 4% PFA for immunofluorescence, and a third was fixed in 10% neutral buffered formalin (NBF) solution (Thermo Fisher Scientific) for histology. In each case, food was carefully removed and the stomach was laid flat on an index card and placed in a cassette with a sponge at top, closed, and fixed in its respective solution. For CFU counts, one-half or one-third stomachs were manually homogenized using a pestle in 0.5 mL of BB10. Serial 10-fold dilutions of stomach homogenate were plated on solid horse blood agar plates containing 4% Columbia agar base (Oxoid, Hampshire, UK), 5% defibrinated horse blood (HemoStat Labs, Dixon, CA) 0.2% β-cyclodextrin (Sigma, St. Louis, MO), 10 μg/mL vancomycin (Sigma), 5 μg/mL cefsulodin (Sigma), 2.5 U/mL polymyxin B (Sigma), 5 μg/mL trimethoprim (Sigma), 8 μg/mL amphotericin B (Sigma), and bacitracin (200 μg/mL) to eliminate normal mouse microbiota growth. Plates were incubated at 37°C using a tri-gas incubator (10% CO_2_, 10% O_2_; Thermo Fisher Scientific) for 4-5 days.

### Competition experiments

Mice were infected with a 1:1 ratio of 10^7^ CFU or wild-type *H. pylori* and the isogenic straight rod mutant (Δcsd6) or the *csd6* complemented strain. After one week, stomachs were removed, divided in half, and plated to determine bacterial loads as CFU/gram of stomach. Wild-type bacterial output was plated on horse blood plates containing the antibiotics described above. The Δ*csd6* mutant was selected on horse blood plates with chloramphenicol (15 μg/mL), and the *csd6* complemented strain was selected on horse blood plates with kanamycin (25 μg/mL).

### Immunofluorescence of thick longitudinal mouse stomach sections

Tissues from mouse stomachs were processed for confocal immunofluorescence microscopy as described in (8, 32), with minor modifications. Gastric tissue was fixed in 4% PFA for 1-2 hours at room temperature. Tissue was embedded in 4% agarose in 1X phosphate-buffered saline (PBS) (pH 7.4) (Gibco) and sectioned using a vibratome to generate 100 – 200 μm thick longitudinal sections that include the limiting ridge at the forestomach/glandular junction to the pyloric junction with the duodenum. Tissue sections were then permeabilized in blocking buffer (3% bovine serum albumin (Sigma Aldrich); 1% Saponin (Sigma Aldrich); 1% Triton X-100 (Sigma Aldrich)) in 1X PBS (pH 7.4) (Gibco). Anti-H. *pylori* rabbit polyclonal antibody (1:1,000 dilution) was used to immunostain *H. pylori* in the tissue overnight at 4°C. The sections were then washed 5X with blocking buffer and incubated with a goat anti-rabbit Alexa Fluor-647 conjugate antibody (1:2,000) to visualize bacteria in tissue (Molecular Probes), and 4’, 6-Diamidino-2-phenylindole (DAPI) (0.1 μg/mL) to stain nuclei for 2 hrs at room temperature. The sections were then washed 5X with blocking buffer and mounted onto standard glass microscope slides with secure imaging spacers (20 mm diameter x 0.12 mm thick, Electron Microscopy Sciences) or hand-made imaging spacers using parafilm (0.12 mm thick). Pro-Long Diamond Antifade medium (Molecular Probes) was added before mounting on coverslips.

### Confocal microscopy

Tissue samples were imaged with a Zeiss LSM 780 NLO confocal and multi-photon microscope with a 40 X oil immersion objective lens (EC Plan-Neofluar 40 X/1.30 oil) and Z-stacks (355 μm (w) x 355 μm (h)) were generated using the ZEN acquisition software program. Images were acquired at a frame size of 1,024 x 1,024 with 8-bit depth and at a frame rate speed of 8 frames per second. Z-stacks were generated with a slice interval of 0.5 μm and penetrated 40-50 μm into the section. For each tissue section, multiple Z-stacks (ranging from 25-30) were acquired to capture the full length of longitudinal sections that include the limiting ridge of the forestomach to the glandular junction to the pyloric junction with the duodenum. For all Z-stacks, a collection of non-overlapping images was acquired by manual translation of the microscope stage.

### Volumetric image analysis and quantitation of H. pylori in the stomach

Quantitation of *H. pylori* within individual gastric glands was performed using the Volocity 3D-image analysis software program, as described in (8, 32), with minor modifications. 3D-reconstructed images were imported onto Volocity and the total volume (μm^3^) for individual bacterial cells was determined. The mean volume for a bacterium (9.5 μm^3^) was used to calculate the total number of bacteria near or at the surface epithelium and within gastric glands. The same measurement protocol was applied across all tissue samples analyzed for wild-type *H. pylori*, the Δ*csd6* mutant, and for *csd6* complemented bacteria. Analysis of 3 sections (>500 μm apart) provided consistent results in bacterial number counts. Our bacterial localization studies included analysis of 1-3 sections per infected mouse. Three sections were analyzed per mouse after one week post-infection and 2-3 sections were examined per mouse at one month post-infection.

### Histologic evaluation of *H. pylori* infected stomachs

Longitudinal gastric strips from the lesser curvature that include the squamocolumnar junction through the proximal duodenum were fixed in 10% NBF. Samples were paraffin embedded, sectioned (5 μm thick), and stained with Hematoxylin and Eosin (H&E) by the Experimental Histopathology Core at the Fred Hutchinson Cancer Research Center. Slides were interpreted and scored using the scoring criteria adapted from (34), shown in Table S2, by a veterinary pathologist (S.E.K.) who was blinded to the experimental details. The individual lesion scores of every mouse in each group were evaluated and compared for inflammation, epithelial defects, oxyntic atrophy, hyperplasia, and metaplasia in the corpus and inflammation and hyperplasia in the antrum. Individual scores (0-4) for each criterion were summed to generate a histological activity index (HAI) score. 9-11 samples per group (mock-infected, wild-type, Δ*csd6*, and *csd6* complemented strain) were evaluated at one and three months post-infection.

### Statistical analyses

We used the Kolmogorov-Smirnov (K-S) statistics tool in CellTool to assay the differences in cell shape morphology, including cell length and side curvature distributions, as described in (16, 17). For CFU data and histopathology scores, comparisons of three groups were performed using a Kruskal–Wallis one-way analysis of variance (ANOVA) followed by Dunn’s multiple test corrections, and pairwise comparisons (competition experiments) were performed with the Mann-Whitney U test using GraphPad Prism 7 (GraphPad Software, La Jolla, CA USA). *P* < 0.05 was considered statistically significant. For the histological activity index (HAI), because mucous metaplasia and hyalinosis may develop spontaneously in mice, these sub-scores were excluded from the calculation of the total HAI.

## Acknowledgments

We thank Dr. Manuel Amieva for the wild-type *H. pylori* PMSS1 strain and antibodies, Dr. Julie Huang for advice in preparing gastric tissue for 3-D confocal microscopy, and Simran Handa (supported by a generous donation from the AT&T Foundation to the FHCRC Summer High School Internship Program) and Ericka Pegues for assistance in generating the *csd6* complemented strain. This work was supported by the US National Institute of Health Grant R01 AI136946 to N.R.S., NIH Ruth L. Kirschstein National Research Service Award (NRSA) F31 AI098424 from the NIAID to L.E.M. This research was supported by the Comparative Medicine, Electron Microscopy, Experimental Histopathology and Scientific Imaging Shared Resource of the Fred Hutch/University of Washington Cancer Consortium (P30 CA015704). The content of this article is solely the responsibility of the authors and does not necessarily represent the official views of the National Cancer Institute and the National Institute of Allergy and Infectious Diseases.

## Author contributions

L.E.M and N.R.S. designed the study. L.E.M V.P.O and C.K.L. performed experiments. L.E.M., V.P.O., C.K.L., and N.R.S. performed data analysis and interpretation. S.E.K. evaluated samples for pathology. L.E.M, V.P.O, S.E.K., and N.R.S. wrote the manuscript.

## Additional information

Supplementary information is available for this paper. Correspondence and requests for materials should be addressed to N.R.S.

## Supporting Tables

**Table S1.**
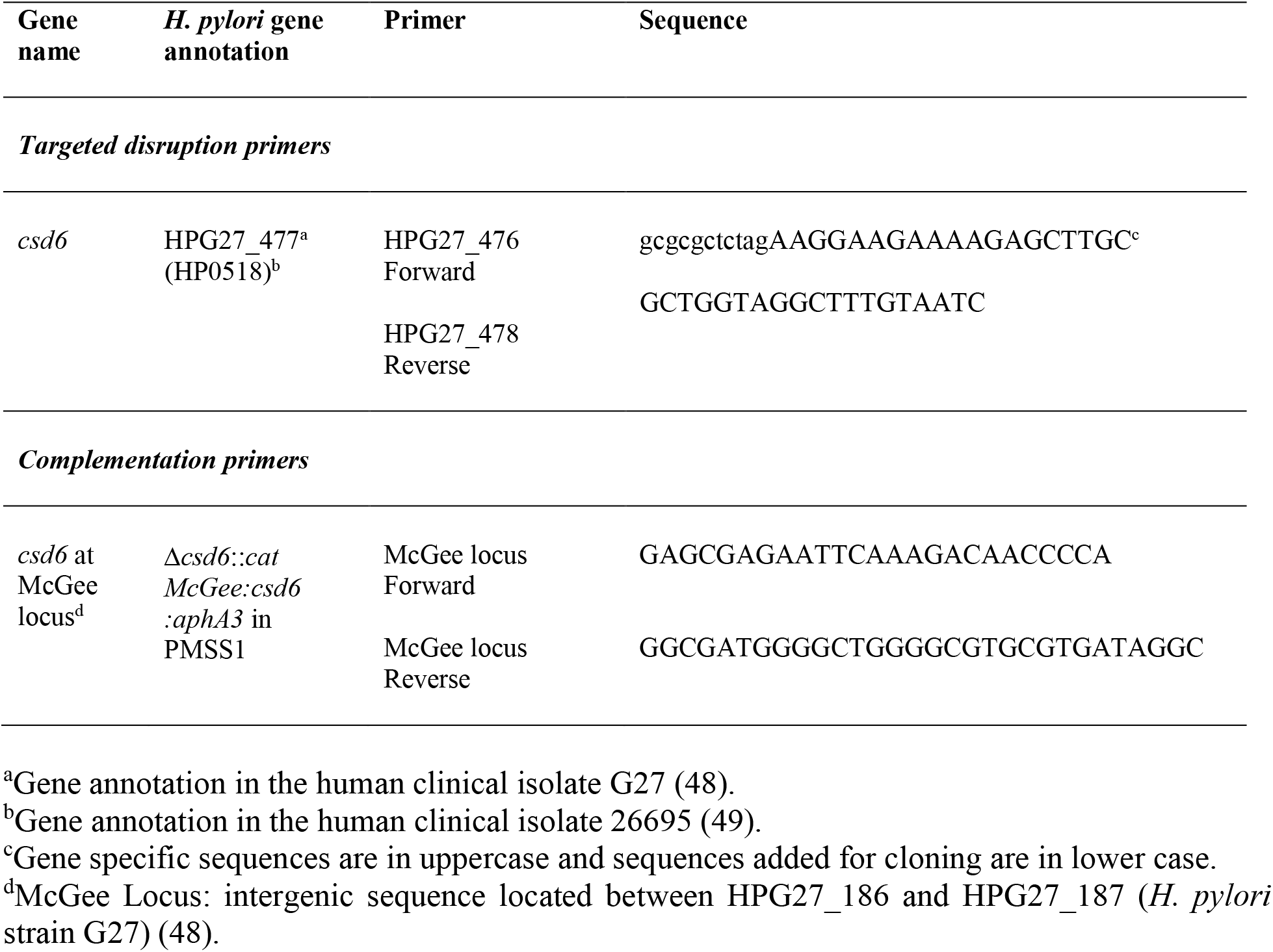
Primers used in this study.

**Table S2.** Histopathologic scoring for inflammation and hyperplasia in mice chronically infected with *H. pylori* for one and three months.

| Score | Inflammation | Hyperplasia | Epithelial defects | Oxyntic atrophy | Metaplasia |
| --- | --- | --- | --- | --- | --- |
| **0** | No inflammation | No hyperplasia | No epithelial defects | No oxyntic atrophy | No metaplasia |
| **1** | Mild patchy or multifocal islands | Mild elongation of mucosa | Tattered epithelial surface | Decreased chief cells; parietal cells intact | < 50% replacement of oxyntic mucosa by antralized glands |
| **2** | Moderate coalescing infiltrate | Increased surface epithelium 2X normal length | Attenuated epithelial surface | Few or no chief cells; parietal cells intact | >50% replacement of oxyntic mucosa by antralized glands |
| **3** | Moderate to severe sheets in mucosa and/or submucosa | Increased surface epithelium 3X normal length | Inapparent surface epithelium | No remaining chief cells; loss of parietal cells | Near complete replacement of glands by antralized mucosa |
| **4** | Severe florid inflammation into muscularis | Increased surface epithelium 4X normal length +/- dysplasia | Mucosal erosions of surface epithelium | No chief cells and few or no parietal cells remaining | Total replacement of glands by antralized mucosa |

## Supplemental Figure Legends

**Figure S1.**
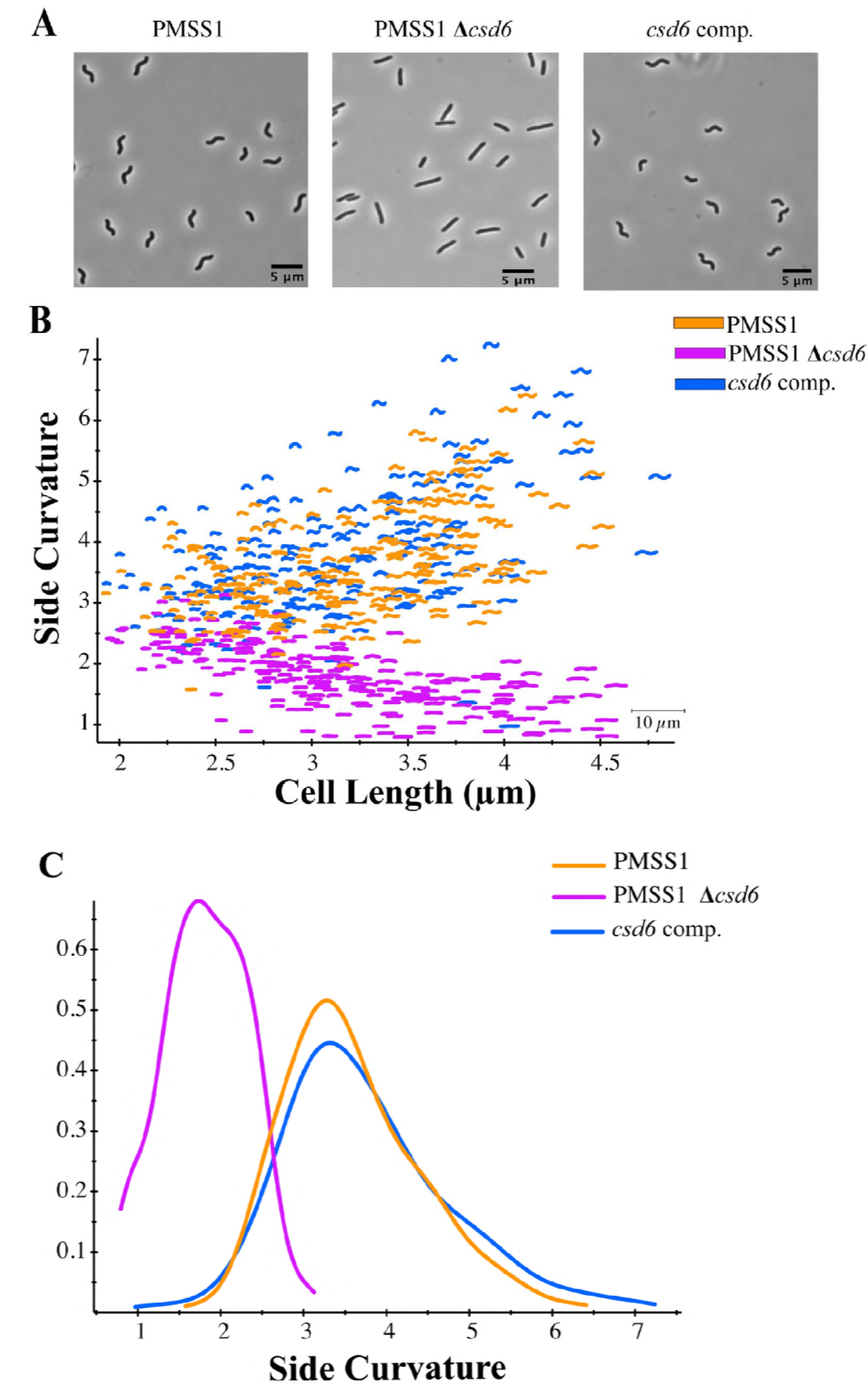
Complementation of *csd6* restores helical cell shape. (A) Representative phase contrast images of wild-type PMSS1 bacteria, straight rod (Δcsd6), and *csd6* complemented bacteria. Images were acquired at 100 X (oil immersion objective). Scale bar = 5 μm. (B) Side curvature vs. cell length (μm) for individual bacterial cells imaged using phase contrast microscopy of wild-type PMSS1 (orange, n=218), Δ*csd6* (magenta, n=230), and the *csd6* complemented strain (blue, n=212). (C) Smooth histograms summarizing the side curvature distributions acquired for each strain shown in B. No significant difference in side curvature distributions were observed between wild-type and the *csd6* complemented strain (p = 0.64078) using Kolmogorov-Smirnov statistics of side curvature distributions. Significant differences in side curvature distributions were observed between wild-type and Δ*csd6*, and between Δ*csd6* and the *csd6* complemented strain, where p<0.00001. Data are from two independent experiments.

**Figure S2.**
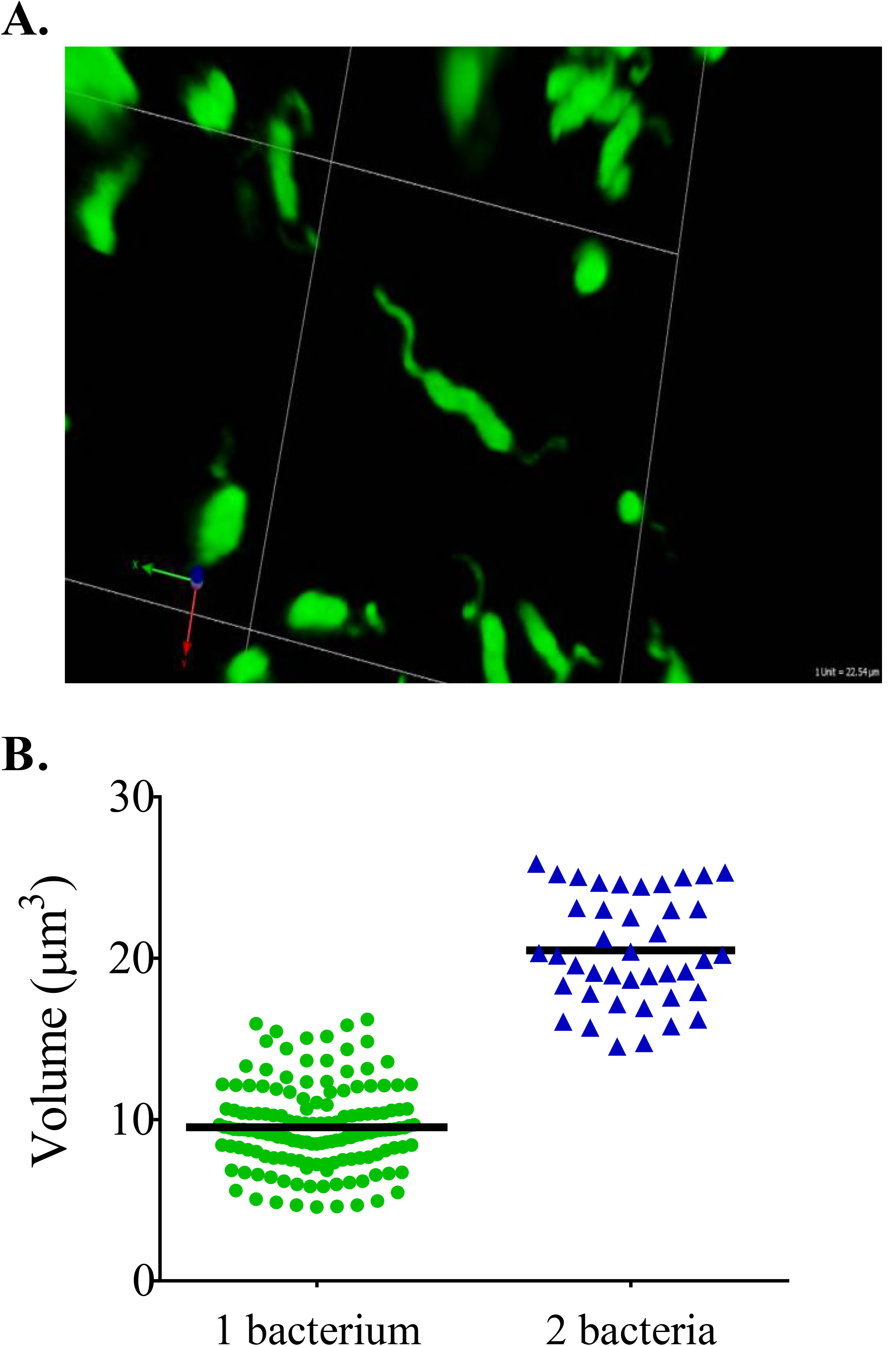
3D-visualization of *H. pylori* and bacterial quantitation by volumetric image analysis. (A) Representative 3D image of wild-type PMSS1 bacteria (green), which was fixed in 2% PFA, embedded in 4% agarose, and sectioned to generate 200 μm thick sections. 3D-images were generated from Z-stacks collected at 63 X (oil-immersion objective) with a Zeiss LSM 780 confocal microscope. (B) Volumetric image analysis of bacterial cells fixed in 2% PFA (n= 203). Bars indicate the mean. Data are from two independent experiments.

**Figure S3.**
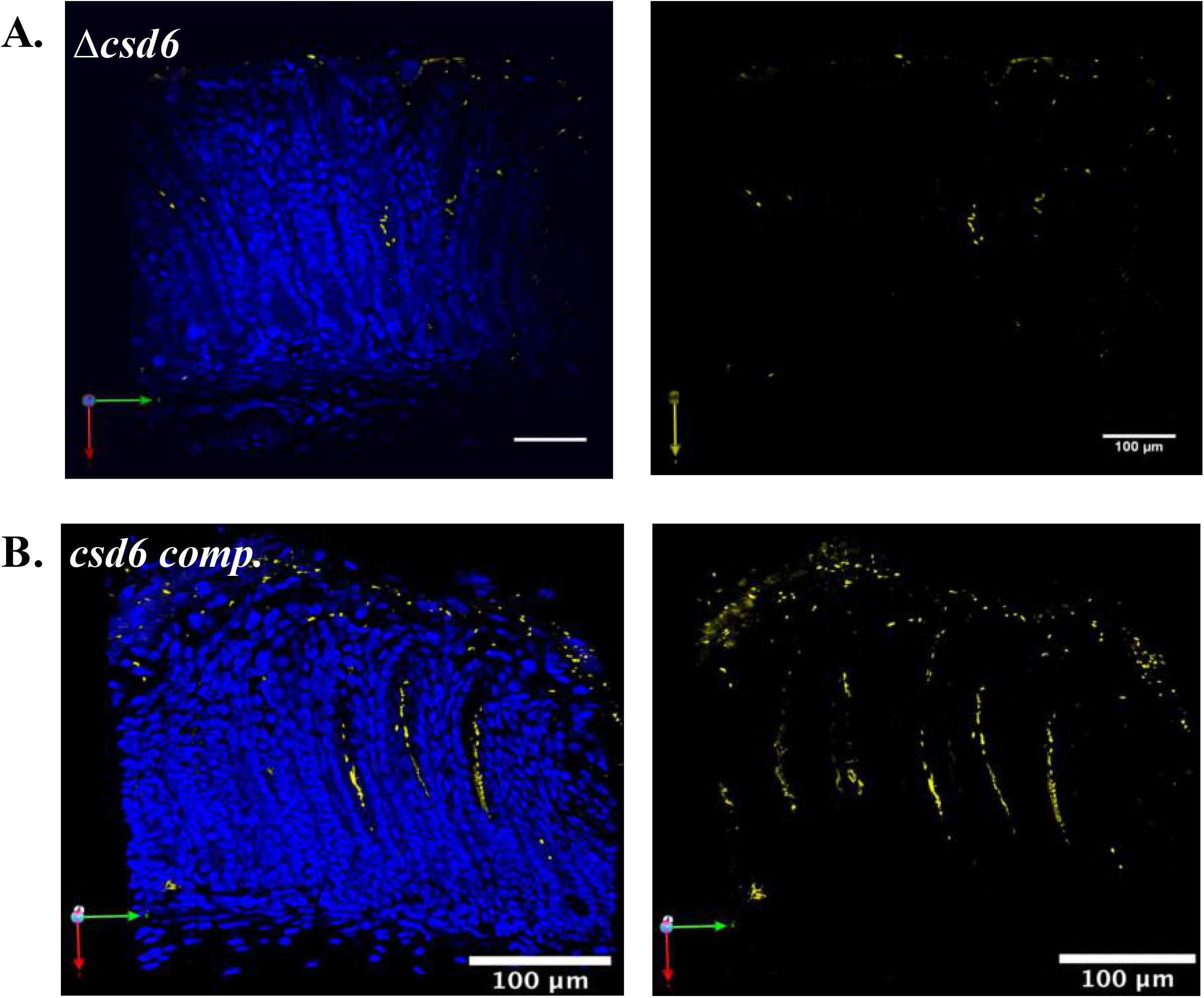
Visualization of bacteria within gastric glands. Thick stomach sections from the one week infections shown in **Figure 2A** were stained for *H. pylori*. Shown are representative images of the antrum of a mouse infected with Δ*csd6* (**A**) or *csd6* complemented (**B**) bacteria. Images are maximum intensity projections of Z-stacks, with blue (DAPI, left panel) staining nuclei and yellow staining *H. pylori*. Scale bars = 100 μm. Volumetric analysis for the mouse in A is found in **Figure 3C** and **B** is in **Figure 3D**.

**Figure S4.**
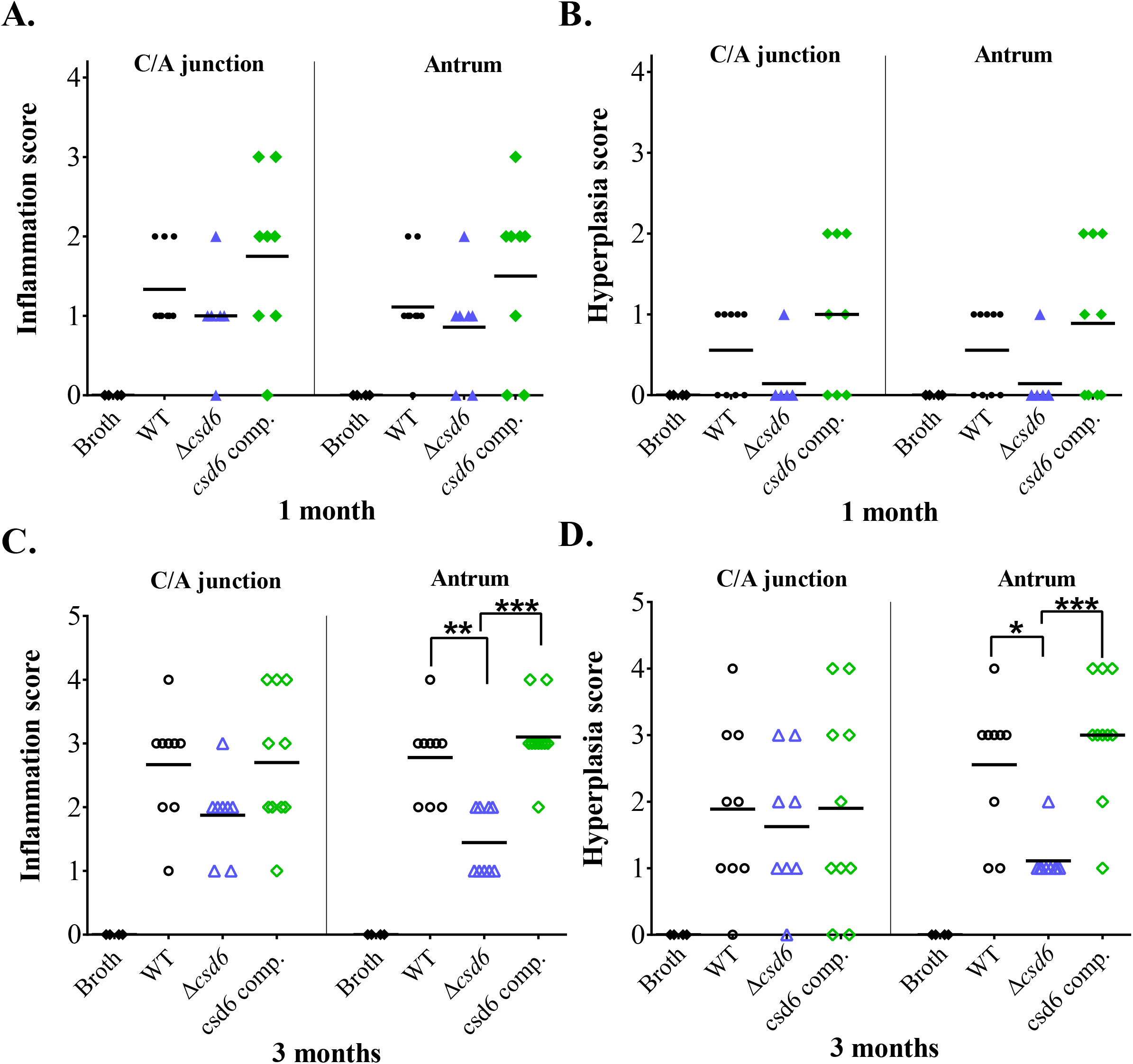
The Δ*csd6* mutant results in decreased inflammation and hyperplasia scores at one and three months of infection. Inflammation (A and C) and hyperplasia (B and D) scores in the corpus/antrum (C/A) junction and antrum at one month (A and B) and three months (C and D) of infection (n=9-11 mice per group). Provided are pathological evaluation scores for all gastric tissue sections analyzed and shown in Fig 5. * *P* <0.05, ** *P* < 0.01, *** *P* < 0.001, Kruskal-Wallis test with Dunn’s multiple test correction.

